# Adaptor protein complex 2 in the orbitofrontal cortex predicts alcohol use disorder

**DOI:** 10.1101/2023.05.28.542637

**Authors:** Patrick J. Mulholland, Stefano Berto, Phillip A. Wilmarth, Christopher McMahan, Lauren E. Ball, John J. Woodward

## Abstract

Alcohol use disorder (AUD) is a life-threatening disease characterized by compulsive drinking, cognitive deficits, and social impairment that continue despite negative consequences. The inability of individuals with AUD to regulate drinking may involve functional deficits in cortical areas that normally balance actions that have aspects of both reward and risk. Among these, the orbitofrontal cortex (OFC) is critically involved in goal-directed behavior and is thought to maintain a representation of reward value that guides decision making. In the present study, we analyzed post-mortem OFC brain samples collected from age- and sex-matched control subjects and those with AUD using proteomics, bioinformatics, machine learning, and reverse genetics approaches. Of the 4,500+ total unique proteins identified in the proteomics screen, there were 47 proteins that differed significantly by sex that were enriched in processes regulating extracellular matrix and axonal structure. Gene ontology enrichment analysis revealed that proteins differentially expressed in AUD cases were involved in synaptic and mitochondrial function, as well as transmembrane transporter activity. Alcohol-sensitive OFC proteins also mapped to abnormal social behaviors and social interactions. Machine learning analysis of the post-mortem OFC proteome revealed dysregulation of presynaptic (e.g., AP2A1) and mitochondrial proteins that predicted the occurrence and severity of AUD. Using a reverse genetics approach to validate a target protein, we found that prefrontal *Ap2a1* expression significantly correlated with voluntary alcohol drinking in male and female genetically diverse mouse strains. Moreover, recombinant inbred strains that inherited the C57BL/6J allele at the *Ap2a1* interval consumed higher amounts of alcohol than those that inherited the DBA/2J allele. Together, these findings highlight the impact of excessive alcohol consumption on the human OFC proteome and identify important cross-species cortical mechanisms and proteins that control drinking in individuals with AUD.

## Introduction

The global burden of alcohol misuse is rising at an alarming rate. In 2019, alcohol use accounted for nearly 2.5 million deaths globally^1^, and the number of alcohol-related deaths between 2019 and 2021 dramatically increased by 38% in the US^2^. Given widespread misuse of alcohol throughout the world, it is not surprising that the prevalence of alcohol use disorder (AUD) among adults is high. For example, lifetime prevalence of AUD in adults is approximately 14% in the US and 23% in Australia with rates considerably higher among males (US: ∼21%; AUS: ∼33%)^3^. Long-term alcohol consumption can lead to an inability to regulate drinking, tolerance to the acute intoxicating effects of alcohol, and physiological and emotional symptoms that appear during withdrawal. The number and severity of these symptoms is used to diagnose individuals with AUD^4^. Characteristics that increase the risk of developing AUD are many and include environmental, cultural, and genetic factors that, in individuals with AUD, each contribute to the loss of control over drinking that often continues in the face of adverse outcomes. This maladaptive behavior is thought to result from alcohol-induced changes in neurocircuits that underlie flexible decision making and risk-reward assessment and includes areas such as prefrontal cortex, ventral and dorsal striatum, and central and basolateral amygdala^5^.

Recent studies from our group and others have focused on the orbitofrontal cortex (OFC) as a critical subregion of the prefrontal cortex that is impacted by acute and chronic exposure to alcohol^6-8^. The OFC is well recognized for its role in goal-directed behavior via its connections with striatal and amygdalar regions and the inputs it receives from sensory stimuli especially that of olfaction and taste that are highly associated with alcohol drinking^9, 10^. The OFC encodes alcohol preference^11^, and deficits in OFC-dependent behaviors are observed in humans diagnosed with AUD^10, 12^, as well as in rodent and non-human primate models of alcohol dependence^10, 13-17^. In humans, these effects are associated with changes in OFC neural excitability under baseline conditions^18^ and during presentations of alcohol related cues^19, 20^; while mice and cynomolgus monkeys with a history of alcohol exposure show alterations in OFC firing and tolerance to the acute inhibitory effects of alcohol^7, 21, 22^. Synaptomic and functional analysis of monkey OFC revealed changes in excitatory glutamatergic synaptic transmission without changes in GABAergic signaling proteins^22^.

We extend our previous findings from rodents and non-human primates by comparing the proteome landscape of the human OFC between age- and sex-matched control cases and individuals with AUD. Because DSM-5 clinical diagnoses were available for AUD cases, we used a supervised machine learning approach to interrogate the relationship between the OFC proteome and diagnosis of AUD and AUD severity. A candidate protein was then validated using bioinformatics and reverse genetics approaches. The results revealed distinct differences in expression of proteins involved in presynaptic glutamatergic activity and mitochondrial metabolism in patient samples that predicted AUD severity and voluntary drinking in mice.

## MATERIALS AND METHODS

### Subjects

Postmortem samples of the lateral OFC (**Fig. 1A**) from 14 pairs of age- and sex-matched samples from controls who abstained or were social drinkers of alcohol (∼5 drinks/week) and those with AUD (**Fig. 1B**), diagnosed according to DSM-5, were analyzed. Lateral OFC was identified^23^ and collected by qualified pathologists under full ethical clearance with informed, written consent from next of kin. Samples were frozen and stored at −80°C until processing. Samples and the demographic information (**Table 1**) were provided by the New South Wales Brain Tissue Resource Centre (Ethnics Committee Approval Number: X11-0107).

**Fig 1.**
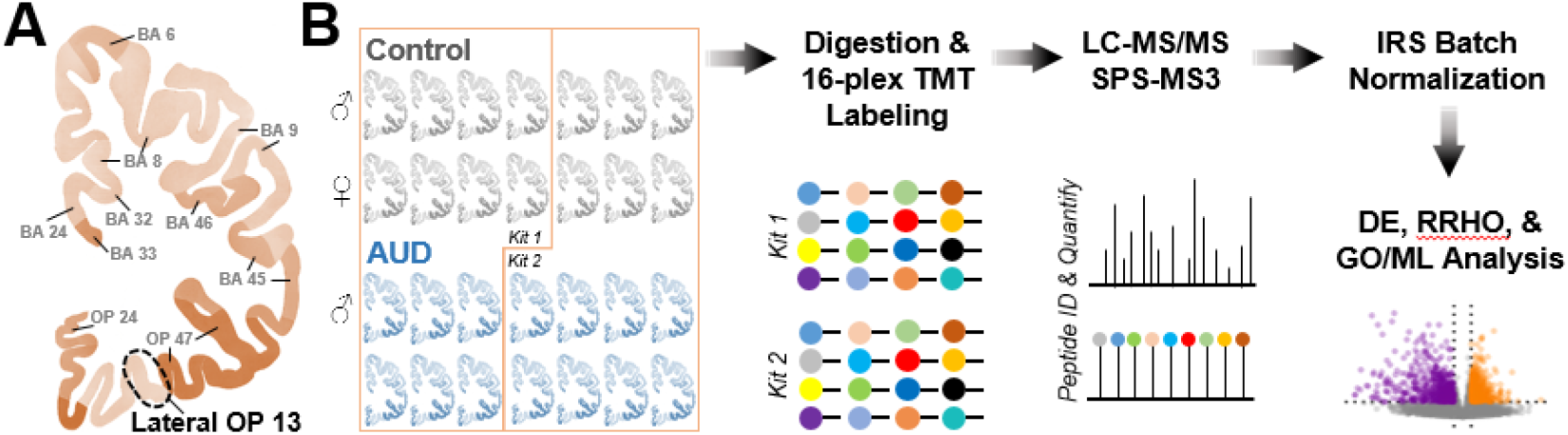
Processing pipeline for proteomics analysis of postmortem samples from the lateral orbitofrontal cortex. **A** Coronal section of human cortex showing Brodmann’s Areas (BA) including the lateral OFC (Orbital Prefrontal (OP) 13) that was acquired for proteomics analysis. **B** Twenty-eight samples from age- and sex-matched controls and AUD cases were digested and labeled using two 16-plex TMT kits before protein identification and quantitation (n = 7 condition/sex). Data were then normalized to control for batch effects across different mass spectrometry runs, and differential expression (DE), rank-rank hypergeometric overlap (RRHO), gene ontology (GO) enrichment, and machine learning (ML) analyses were completed.

**Table 1.**
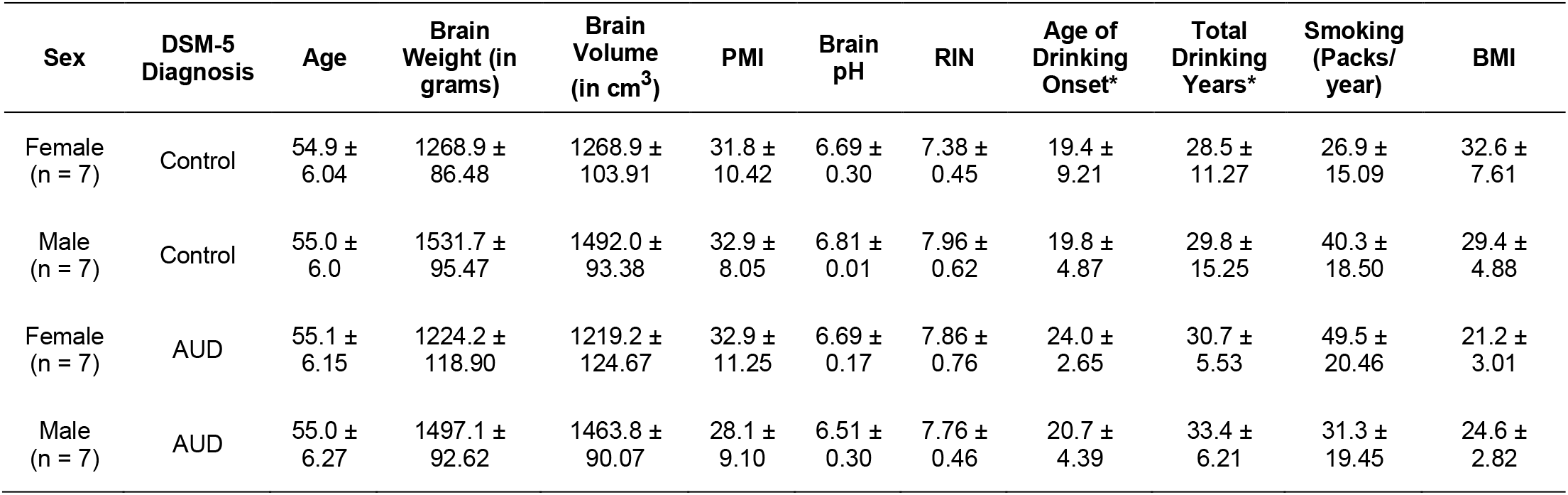
Summary of the demographics of control subjects and subjects with Alcohol Use Disorder. Data are reported as mean and standard deviation. *Denotes estimated value where an age of onset was recorded as 25 years of age in unknown cases (see^98^). RIN, RNA integrity number; PMI, postmortem interval; BMI, body mass index. Mean ± s.e.m.

| Sex | DSM-5 Diagnosis | Age | Brain Weight (in grams) | Brain Volume (in cm<sup>3</sup>) | PMI | Brain pH | RIN | Age of Drinking Onset* | Total Drinking Years* | Smoking (Packs/year) | BMI |
| --- | --- | --- | --- | --- | --- | --- | --- | --- | --- | --- | --- |
| Female (n = 7) | Control | 54.9 ± 6.04 | 1268.9 ± 86.48 | 1268.9 ± 103.91 | 31.8 ± 10.42 | 6.69 ± 0.30 | 7.38 ± 0.45 | 19.4 ± 9.21 | 28.5 ± 11.27 | 26.9 ± 15.09 | 32.6 ± 7.61 |
| Male (n = 7) | Control | 55.0 ± 6.0 | 1531.7 ± 95.47 | 1492.0 ± 93.38 | 32.9 ± 8.05 | 6.81 ± 0.01 | 7.96 ± 0.62 | 19.8 ± 4.87 | 29.8 ± 15.25 | 40.3 ± 18.50 | 29.4 ± 4.88 |
| Female (n = 7) | AUD | 55.1 ± 6.15 | 1224.2 ± 118.90 | 1219.2 ± 124.67 | 32.9 ± 11.25 | 6.69 ± 0.17 | 7.86 ± 0.76 | 24.0 ± 2.65 | 30.7 ± 5.53 | 49.5 ± 20.46 | 21.2 ± 3.01 |
| Male (n = 7) | AUD | 55.0 ± 6.27 | 1497.1 ± 92.62 | 1463.8 ± 90.07 | 28.1 ± 9.10 | 6.51 ± 0.30 | 7.76 ± 0.46 | 20.7 ± 4.39 | 33.4 ± 6.21 | 31.3 ± 19.45 | 24.6 ± 2.82 |

### TMT labeling and fractionation of peptides

Tissue sample were solubilized in 9M Urea (Sequanal grade, ThermoFisher), 50mM HEPES (pH 8.0), 100units/mL universal nuclease (ThermoFisher) with brief sonication. Homogenates were centrifuged at 10,000 x *g* for 5 minutes at room temperature to remove debris and the concentration of protein in the supernatant determined by Pierce BCA assay (ThermoFisher). Protein (180μg/sample) was reduced in 5mM dithiothreitol for 30 minutes at 37°C and cysteines alkylated with 10mM iodoacetamide for 45 minutes at 25°C in the dark. The concentration of urea was diluted to less than 2M with 50mM HEPES (pH 8.0). Lys-C endoprotease (Waco) digestion was performed at an enzyme:protein ratio of 1:100 for 2 hours at 25°C followed by trypsin (Sigma) digestion at an enzyme:protein ratio of 1:50 overnight at 37°C. The digestion reactions were quenched by acidifying with 5% (v/v) formic acid. Digested peptides were desalted using 1 cc (50mg) Sep-Pak Vac tC18 solid phase extraction cartridges (Waters) and the concentration determined using the Pierce quantitative colorimetric peptide assay (ThermoFisher). Peptides were dried by vacuum centrifugation.

Two tandem mass tag (TMT) kits (16-Plex TMTpro, ThermoFisher) were used for isobaric labeling^24^. Samples were pseudo-randomly assigned to each kit balanced for age, sex, and condition, and pairs of age- and sex-matched samples were processed within the same kit. Digested peptides (100μg per TMT channel) were resuspended in 100μL of 200mM HEPES (pH 8.5). TMT reagents were dissolved in 20μL of anhydrous acetonitrile and added to the peptide samples. Reactions were incubated for 2 hours at room temperature in the dark and stored at -80°C while confirming the labeling efficiency. For normalization and batch correction across the two TMT experiments, a pooled reference standard comprised of 27μg of protein from each sample was labeled with the 126 or 134N reagent. To check for sufficient labeling and accurate mixing, 2μL from each reaction were quenched, combined, and analyzed by LC-MS/MS as described below. Mass spectrometric data were searched against a UniProt reviewed human protein database (20,354 entries downloaded 02/04/2020) using the SequestHT algorithm in Proteome Discoverer 2.5. TMTpro tags on the peptide N-termini and lysines were included as variable modifications. Percentage of peptide spectral matches (PSM) with labeled N-termini and/or lysine as compared to the total number of PSMs identified was >95%. To assess the accuracy in combining the samples at equal amounts, the data were searched with the TMTpro labels as fixed modifications and the sum of reporter ion intensities for each reagent channel were compared. After confirming efficient labeling and equal mixing ratios, the TMT reactions were quenched with 5% hydroxylamine, combined, and dried by vacuum centrifugation. Labeled peptides were desalted using 3cc (200mg) Sep-Pak Vac tC18 cartridges (Waters). A 100-μg aliquot of the labeled, combined, desalted peptides was separated into 9 fractions with the high pH reversed-phase peptide fractionation kit (ThermoFisher). Fractions with increasing percentages of acetonitrile (ACN) in 0.1% triethylamine as well as the flow through were collected. Elution solutions contained 5%, 7.5%, 10%, 12.5%, 15%, 17.5%, 20%, and 50% ACN. Eluted peptides were dried and desalted using Stage tips.

### LC-MS/MS data acquisition

Multiplexed, fractionated, TMT labeled peptides were analyzed by LC-MS/MS using an EASY nLC 1200 in-line with the Orbitrap Lumos Tribrid mass spectrometer (ThermoFisher). Peptides were pressure loaded at a maximum of 1,000 bar, and separated on a C18 reversed phase column (Acclaim PepMap RSLC, 75μm x 50cm (C18, 2μm, 100 Å)), ThermoFisher)) with a gradient of 5% to 35% solvent B over 180 min (Solvent A: 2% ACN, 0.1% formic acid; Solvent B: 80% ACN, 0.1% formic acid) at 300nL/min. The column was thermostated at 45°C.

To reduce the impact of co-isolated ions on reporter ion intensities, data were acquired using synchronous precursor selection (SPS-MS3)^25^. MS1 scans were detected in the Orbitrap analyzer at a resolution of 120K. For MS2 scans, precursors were isolated with a 0.7Da window, an AGC target of 1×10^4^, and automatic maximum injection time. Precursors were fragmented by collision induced dissociation with a normalized collision energy (NCE) of 30% and analyzed in the ion trap. Ions within a 10-ppm window of a previously analyzed precursor were dynamically excluded for 15 sec. SPS-MS3 scans were acquired on the top 10 most intense fragment ions from the MS2 scan. MS3 precursors were isolated using a 2 Da window and fragmented by high energy collisional dissociation at a NCE of 55% with an AGC of 1×10^5^ and maximum injection time of 120ms. MS3 fragment ions were analyzed in the Orbitrap at 60K resolution within a mass range of 110-500m/z.

### Database searching, peptide identification, quantitation, and functional annotation

The Comet search engine (v2016.03)^26^ was used: 1.25Da monoisotopic peptide mass tolerance, 1.0005Da monoisotopic fragment ion tolerance, fully tryptic cleavage with up to two missed cleavages, variable oxidation of methionine residues, static alkylation of cysteines, and static modifications for TMTpro labels (at peptide N-termini and at lysine residues). The protein sequence collection searched was canonical human Swiss-Prot FASTA sequences downloaded 11/30/2021 from UniProt.org. Common contaminants (175 sequences) were added and sequence-reversed entries were concatenated for a final protein FASTA file of 41,100 sequences. Each TMT plex had 9 fractions for a total of 18 instrument files that were processed with the PAW pipeline^27^. Binary files were converted to text files using MSConvert^28^. Python scripts extracted TMTpro reporter ion peak heights and fragment ion spectra in MS2 format^29^.

Top-scoring PSMs were filtered to a 1% FDR using an interactive GUI application to set thresholds in delta-mass histograms and conditional Peptide-prophet-like linear discriminant function^30^ score histograms where incorrect delta-mass and score histogram distributions were estimated using the target/decoy method^31^. Filtered PSMs were assembled into protein lists using basic and extended parsimony principles and required two distinct peptides per protein per plex. The final list of identified proteins, protein groups (indistinguishable peptide sets), and protein families (highly homologous peptide sets) were used to define unique and shared peptides for quantitative use. Total (summed) reporter ion intensities were computed from PSMs associated with all unique peptides (final grouped protein context) for each protein. Intensities of duplicate pooled internal standards for each protein were averaged in each TMT plex to create local protein intensity scales. Two local scales were adjusted to a common scale (the geometric mean of the local scales) and scaling factors applied to TMT intensities of the 14 biological samples in each plex. This internal reference scaling^32^ puts all 28 biological sample protein intensities on a common scale. Mass spectrometry data have been deposited to the ProteomeXchange Consortium via the PRIDE partner repository^33^ with the dataset identifier PXD041727.

### Machine learning for AUD separation

To determine if proteins can separate control and AUD cases, as well as identify proteins that can separate the severity of AUD diagnosis (mild, moderate, or severe AUD), we applied sparse partial least squares discriminant analysis (sPLS-DA) to the proteomics data set, which was implemented using the mixOmics package in R. sPLS-DA is a supervised machine learning algorithm that performs variable selection and classification in one operation and uses a sparsity assumption to identify a small number of important variables for separation^34^. Five-fold repeated cross validation with 50 repetitions was applied to the model. Separation of control and AUD cases was determined by component 1 and 2 projections, and loading coefficients for top weighted variables were reported.

### Bioinformatics and reverse genetics

Two additional open-source bioinformatics analyses were performed following our previous methods^22, 35-39^. GeneWeaver software system, a database containing major curated repositories and functional genomics results obtained from experiments across 9 species^40^, was queried to retrieve data that implicate the gene that encodes AP2α1 in alcohol-related phenomena. Next, we performed targeted analyses using existing genetic and phenotypic data in GeneNetwork^41, 42^. We correlated prefrontal *Ap2a1* expression (probe set: ENSMUSG00000060279) in adult male and female alcohol-naïve BXD recombinant inbred strains of mice (GeneNetwork data set: DOD_BXD_PFC_GWI_CTL_RNA-Seq_ComB_(Dec19)_TPM_Log2) with published phenotypic drinking data from control and CIE-exposed BXD strains^43^. Next, BXD strains were genotyped at the *Ap2a1* interval to assess the contribution of the *B* or *D* allele in alcohol drinking behavior.

### Statistics

Sample size was estimated by power analysis using G*Power^44^. Protein intensity values for each biological sample were compared for differential protein expression using two factor [sex × condition] analysis. Differential protein intensity analysis was performed in R using a linear model as following: *lm(protein intensity ∼ Condition + Sex + SVs + Condition*Sex)*. Surrogates Variables (SVs) were calculated using *SVA* (v3.42) in R based on a “2-step” method with 100 iterations^45^. A total of two SVs were included in the analysis. Post-hoc analysis was performed in R using *emmeans* package (v1.8.4). Resultant p-values were corrected by Benjamini-Hochberg procedure. FDR < 0.05 was used to determine statistical significance. Functional annotation of the differentially expressed proteins (DEPs) was performed in ToppGene^46^ using the full list of identified proteins as the background set. Rank-rank hypergeometric overlap (RRHO2)^47^ was used to assess concordance of DEPs in males and females with AUD. Drinking data were examined for normal distributions and equality of variance using Q-Q plots, and drinking data in BXDs were lognormally distributed. Four-factor linear mixed models of log-transformed values were calculated in MATLAB (R2022a) with α set to 0.05.

## RESULTS

### Drinking characteristics

As shown in **Table 1**, estimated total drinking years were similar between control and AUD cases (∼28.5–33 years), but the number of drinks per week were markedly different. In control cases, alcohol drinks/week ranged from 3.2–7.1 (5.3–10.3 grams of alcohol/day), whereas AUD individuals consumed 96.0–146.9 alcohol drinks each week (199.1–256.4 grams of alcohol/day; **Fig. 2A**). Of the individuals with AUD, 5 met criteria for mild AUD, 2 met criteria for moderate AUD, and 7 met criteria for severe AUD, according to DSM-5 diagnostic criteria.

**Fig 2.**
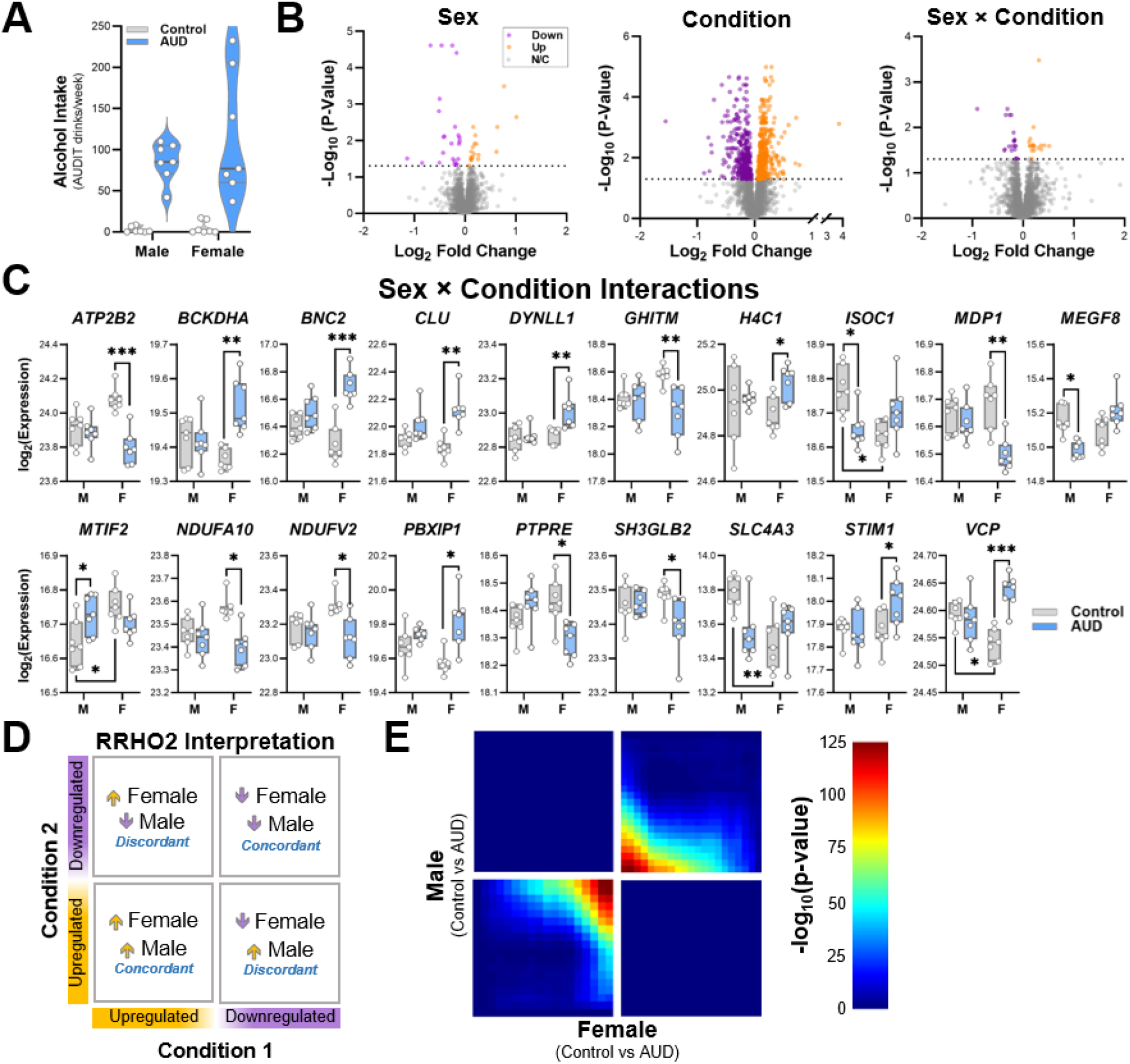
Proteomic adaptations in the OFC across sex and in AUD cases. **A** Drinks per week in controls and individuals with AUD. n = 7 condition/sex. Lines denote median plus interquartile range. **B** Volcano plots showing differentially expressed proteins (DEPs) in the lateral orbitofrontal cortex for each factor (sex, condition) and the interaction between sex and condition among controls and AUD cases. **C** Post-hoc analysis of proteins with significant sex by condition interactions in male and female control and AUD cases. *p < 0.05. ** p < 0.01, *** p < 0.001; n = 7 condition/sex. Box plots show median and interquartile range. **D.E** Interpretation of RRHO2 plots and expression concordance between males and females with AUD.

### Proteomics analysis of human lateral OFC

Age- and sex-matched pairs of lateral OFC (**Fig. 1A**) samples were pseudo-randomly divided across two TMTpro experiments (**Fig. 1B**). Following peptide identification and quantification for the individual kits, batch effects from separate MS runs were removed using IRS normalization^32^ (**Supplemental Fig. 1**). Then, differential expression, concordance, Gene Ontology enrichment, and machine learning analyses were performed (**Fig. 1C**). There were 754,217 acquired MS2/MS3 scans with 266,029 scans passing the 1% FDR filtering. PSMs mapped to 4,957 protein groups, with 4,851 identified protein groups after excluding decoy and common contaminant matches. The final number of quantifiable proteins observed in both TMT plexes was 4,503, and data were analyzed using two-factor analysis [sex × condition]. Volcano plots for each factor and the interaction between sex and condition are shown in **Fig. 2B**. Analyses revealed 43 DEPs between males and females (main effect of sex) and 39 proteins that met criteria for a significant sex × condition interaction. Analyses also revealed major differences in the OFC proteome of individuals with AUD (main effect of condition). Because there were proteins with significant interactions, we performed post-hoc analyses to determine if AUD differentially affected these proteins in males and females. Of the 39 proteins with significant interactions, posthoc tests reached statistical significance for 19 proteins (**Fig. 2C**). Four proteins were significantly different between male and female controls (ISOC1, MTIF2, SLC4A3, and VCP), 3 were significantly different between male controls and males with AUD (ISOC1, MEGF8, and MTIF2), and 15 proteins were significantly different between female controls and females with AUD. RRHO2 analysis revealed high expression concordance for DEPs in males and females with AUD (**Fig 2D**, Fisher’s exact test, *p* < 0.0001).

### Gene ontology analysis

Next, we completed Gene Ontology (GO) enrichment analysis to determine processes and functions that were disrupted by sex and AUD. We first performed GO enrichment analysis on the complete list of 4503 unique proteins comparing them with the full human transcriptome (**Fig. 3A**). Overall, the full list of proteins identified in our screen were enriched in ‘cadherin,’ ‘cell adhesion’, and ‘cytoskeletal protein binding’. These proteins showed enriched expression in ‘glutamatergic synapses’ (pre- and post-synaptic compartments) and functions that ‘regulate intracellular’ and ‘membrane protein transport’. Because the differential expression analysis revealed that AUD differentially affects proteins in males and females (see **Fig. 2C** for interactions), we performed separate GO enrichment analysis for each sex comparing the DEPs to all proteins identified in our screen. Comparing the 47 proteins that differed between males and females, DEPs were enriched in ‘extracellular matrix’ (ECM), ‘basement membrane’, and ‘integrin signaling’, as well as ‘processes regulating axon guidance’ and ‘axonogenesis’ (**Fig. 3B**). Although there were minor differences, GO enrichment analysis on AUD-dependent protein changes was similar for control vs AUD in males (**Fig. 3C**) and females (**Fig. 3D**), which showed proteins involved in ‘disruption of mitochondrial functioning’ and ‘transmembrane transporter activity’. Additional analysis revealed that down-regulated proteins were enriched in catabolic and metabolic processes within mitochondria, whereas up-regulated proteins were enriched at glutamatergic synapses and regulate synaptic vesicle cycling and release. Finally, GO analysis indicated that DEPs in males and females with AUD were enriched in phenotypes related to general categories for nonverbal communication and social behaviors (**Fig. 3E**). DEPs in individuals with AUD were also enriched in phenotypes for ‘seizure’ and ‘abnormal cerebral morphology’, all of which were not identified when analyzing the full list of identified proteins.

**Fig 3.**
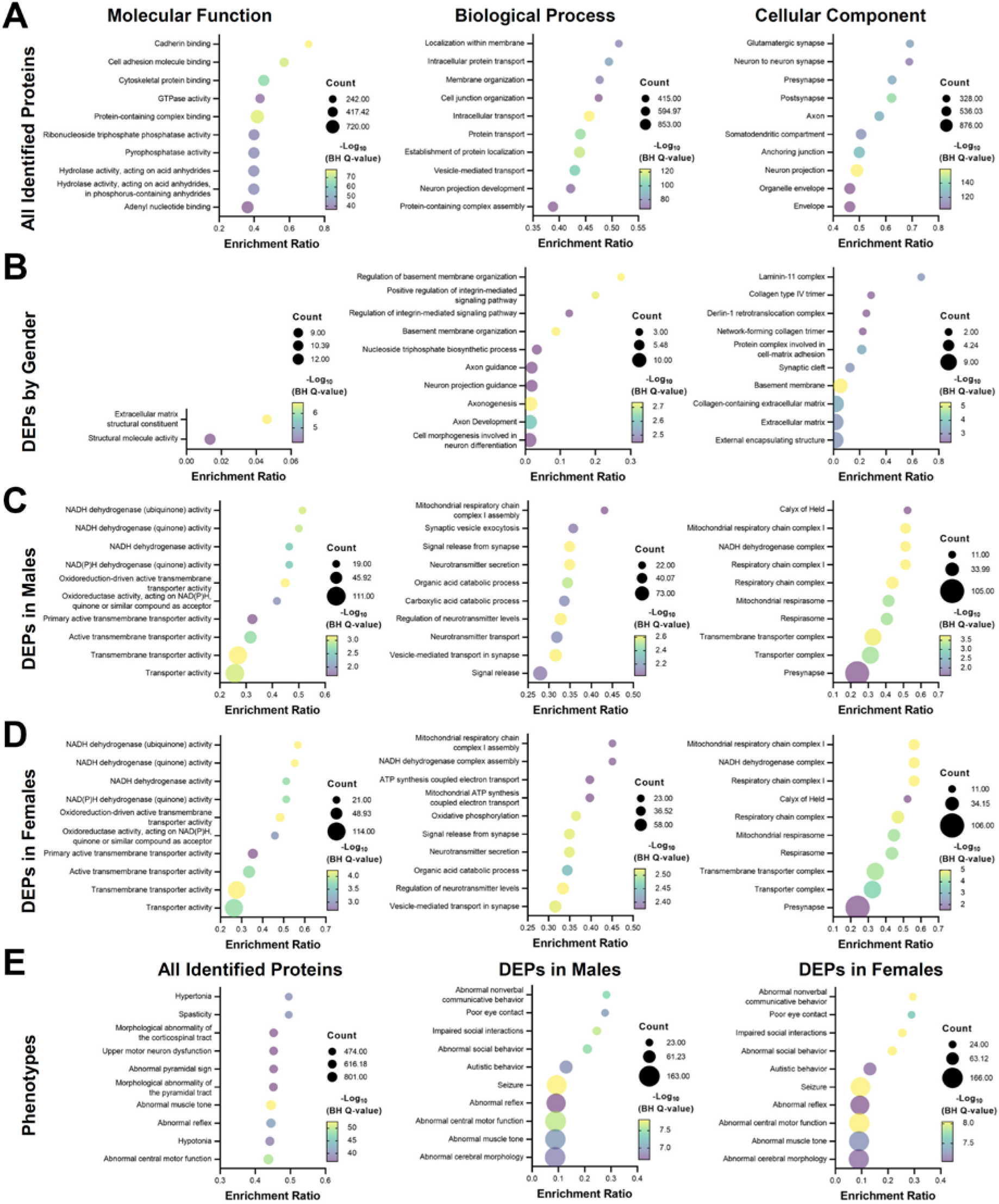
Gene ontology (GO) enrichment analysis - molecular function, biological process, and cellular component - of the OFC proteome in AUD. **A** GO enrichment analysis of all 4503 identified proteins compared with the full human transcriptome. **B** GO analysis of the DEPs when comparing gender. **C** Enrichment analysis of DEPs in male control and AUD cases. **D** GO enrichment analysis of DEPs in female control and AUD cases. **E** Phenotype analysis of gene ontology enrichment in all identified proteins and DEPs in male and female control and AUD cases.

### AUD prediction

As a final analysis, we applied a supervised machine learning approach (i.e., sPLS-DA) to identify proteins that best distinguish between controls and AUD samples and proteins that predict AUD severity (mild, moderate, and severe AUD diagnosis). There was clear discrimination between control and AUD samples (**Fig. 4A**), and the proteins with the top 50 loading coefficients (weights) for the first component of the model are shown in **Fig. 4B**. Interestingly, proteins with positive loadings in the discrimination model had different molecular and biological functions and localized to distinct compartments of cells compared with proteins with negative loadings (**Fig. 4C**). Whereas proteins with positive loadings were generally involved in metabolic processes within mitochondria, proteins with negative loadings regulate transmembrane transporter activity and vesicle endocytosis (e.g., AP2A1 encoding the adaptor protein complex 2 subunit α1) and release within glutamatergic synapses. Similarly, sPLS-DA separated samples based on the severity of AUD diagnosis (**Supplemental Fig. 2A**). Control samples were distinguished from those obtained from both mild and severe AUD cases. The two samples from moderate AUD cases clustered with the severe AUD samples. Loadings coefficients for severity discrimination analysis are shown in **Supplemental Fig. 2B**. GO enrichment analysis revealed that the 10 proteins in the first component of the model are involved in metabolic and catabolic processes and are largely expressed within mitochondria (**Supplemental Fig. 2C**).

**Fig 4.**
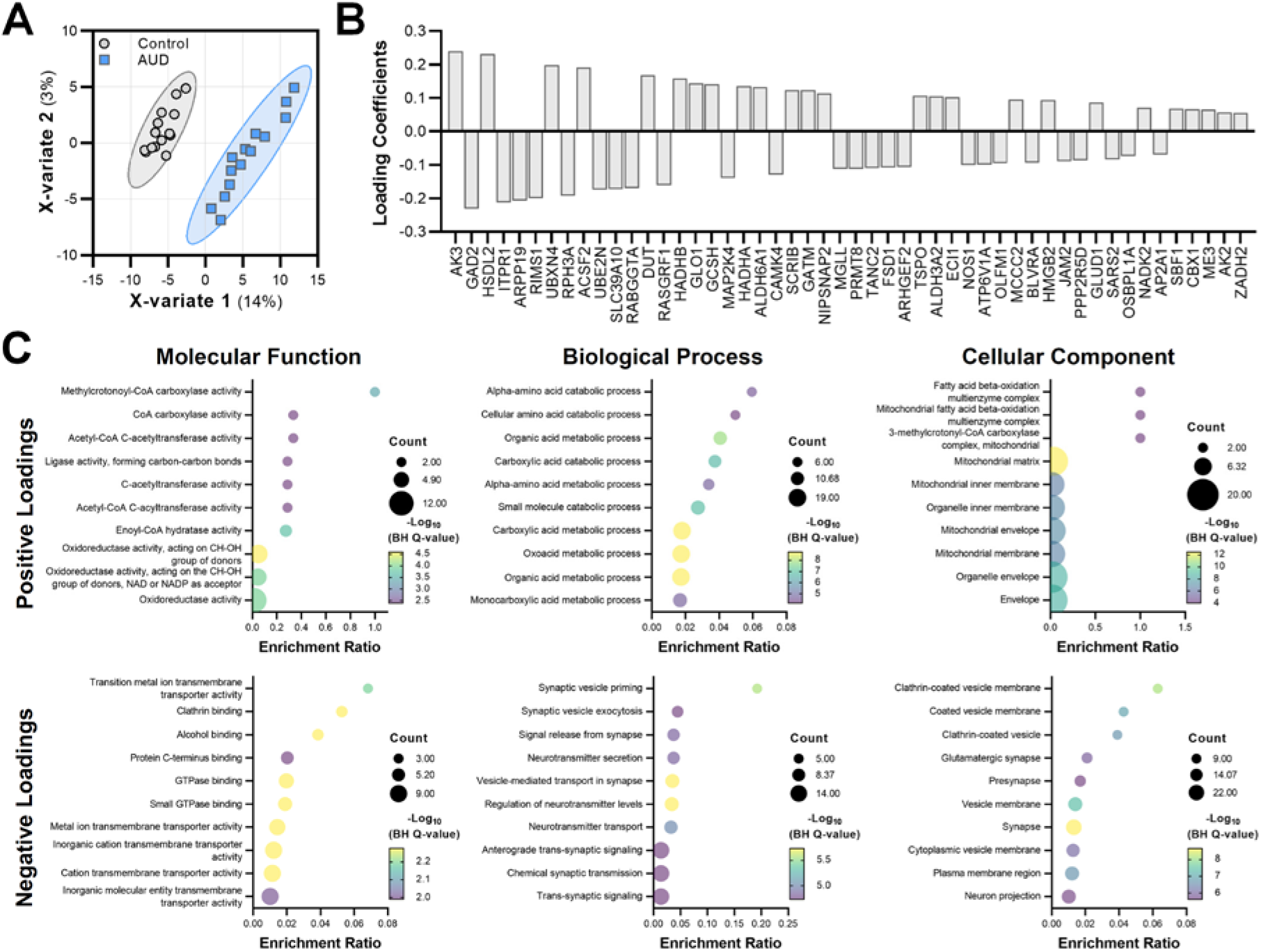
Proteins that separate control and AUD cases. **A** Separation of control and AUD cases **B** Top 50 loading coefficients for the first component of the sPLS-DA model. Bar length indicated importance of each protein to the model. **C** Gene ontology enrichment analysis for positive and negative loading coefficients of proteins that separate control and AUD cases. **D** Bioinformatics analysis linking AP2A1 to alcohol-related behaviors in mouse, monkey, and human. **E** Cortical expression of Ap2a1 correlated with alcohol drinking in 23 control and alcohol dependent BXD strains. F The B allele at Ap2a1 increased alcohol drinking in 44 BXD strains.

### Cross-species validation of AP2A1

In addition to showing differential expression and separating control from AUD cases in this study, machine learning analysis also revealed that AP-2 α1 separated low and heavy alcohol drinkers in our non-human primate study^22^. In addition, we previously reported that expression of AP-2 α1 was reduced in the accumbens of alcohol dependent mice^48^. Because of these relationships, we expanded our analysis on *AP2A1* and alcohol-related behaviors across species. Using a bioinformatics approach, we found that *Ap2a1* was present in QTLs for alcohol preference, conditioned taste aversion, and hypothermia in mice, and brain *AP2A1* transcript expression was altered in individuals with AUD and after acute and chronic alcohol exposure in mice (**Fig. 5A**). A single nucleotide polymorphism (rs1001281) was identified in *AP2A1* that associated with ‘MaxDrinks’^49^, a heritable trait and risk factor for developing AUD^50-52^. Next, we experimentally tested the association between *Ap2a1* and voluntary alcohol drinking in control and alcohol dependent BXD recombinant inbred strains of mice. BXDs are generated by crossing alcohol-avoiding DBA/2J (*D*) and alcohol-preferring C57BL/6J (*B*) mice and are a valuable tool for identifying genetics drivers of variation in alcohol-related behaviors^41^, including drinking^22, 35-39^. In prefrontal cortex, we found that *Ap2a1* expression significantly correlated with voluntary alcohol drinking in control and chronic intermittent ethanol (CIE) exposed BXDs (Main effect of genotype: F_1,40.274_ = 24.655, p < 0.001, *n* = 19 strains; **Fig. 5B**). Using a reverse genetics approach^22, 53^, we compared alcohol intake directly linked to the *Ap2a1* interval in BXDs that inherited the *D* or *B* allele. As shown in **Fig. 5C**, the strains that inherited the *B* allele at *Ap2a1* consumed significantly more alcohol than strains that inherited the *D* allele (Main effect of genotype: F_1,94.59_ = 5.80, *p* = 0.0179; *n* = 44 strains). Together, these results demonstrate that variation in cortical AP2A1 expression regulates alcohol drinking across mice, monkeys, and humans.

**Fig 5.**
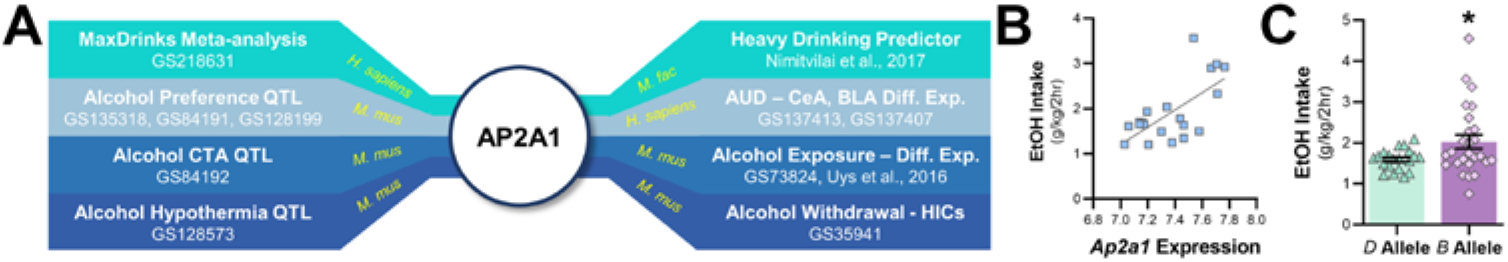
Adaptor protein complex 2 a1 and alcohol-related phenotypes across species. **A** Bioinformatics analysis linking AP2A1 to alcohol-related behaviors in mouse, monkey, and human. CeA, central nucleus of the amygdala; CTA, conditioned taste aversion; Diff. Exp, differential expression; GS, GeneWeaver.org gene set number; HICs, handling induced convulsions; QTL, quantitative trait locus. **B** Cortical expression of Ap2a1 correlated with alcohol drinking in 23 control and alcohol dependent BXD recombinant inbred strains. **C** The B allele at Ap2a1 increased alcohol drinking in 44 BXD strains (*p < 0.05).

## DISCUSSION

The present study extended our findings in mouse and non-human primate models for the study of AUD by interrogating the human OFC proteome in deceased healthy controls and individuals with AUD. First, we found sexual dimorphism in OFC proteins that regulate axons, brain ECM structure, and ECM-cell and ECM-ECM signaling. Our results also provide evidence of mitochondrial, ion channel, and synaptic adaptations in AUD cases that were largely similar in males and females. Using a supervised machine learning analysis, we identified mitochondrial metabolic and presynaptic proteins as predictors of AUD, as well as predictors of AUD severity. In addition, a subset of the differently expression proteins mapped to abnormal social behaviors and social interactions. Finally, we validated an intriguing target using bioinformatics and reverse genetics approaches to demonstrate cross-species control of cortical AP2A1 over alcohol drinking.

### Alcohol-sensitive OFC proteins

Comparison of the OFC proteome identified alcohol-sensitive proteins that are expressed in two major cellular compartments – mitochondria and glutamatergic synapses – and regulate transmembrane transporter activity. Mitochondrial proteins differentially expressed in AUD cases are key components of respiratory chain supercomplexes I, III, and IV that comprise the respirasome, which controls oxidative phosphorylation and generates the mitochondrial electrochemical gradient^54^. The finding of alcohol-responsive mitochondrial proteins replicated previous human cortex postmortem^55-58^ and preclinical^59-61^ studies, which reported changes in mitochondrial metabolic genes and mitochondrial function. The current results demonstrating mitochondrial dysregulation in human OFC are consistent with our proteomic study in the OFC of alcohol drinking macaques^22^, but extended the findings to demonstrate that AUD affects the entire respirasome supercomplex. Individuals with AUD exhibit OFC degeneration^62, 63^ and deficits in cognitive control, reward-based decision-making, and behavioral flexibility, all of which depend on an intact OFC^9, 12^. Since the brain requires high energy demands to maintain proper function and mitochondria can regulate firing rate set points^64, 65^, mitochondrial deficits in AUD cases may have profound effects on overall OFC function that could contribute to oxidative damage, cellular pathology, and impairments in OFC-dependent cognitive, behavioral, and social functioning.

A major finding was the identification of alcohol-sensitive proteins that regulate cellular firing, passive and active biophysical membrane properties, and synaptic function. The dysregulated proteins that control biophysical membrane properties include subunits of the Na^+^/K^+^-ATPase responsible for maintaining the electrochemical gradient (ATP1B1, ATP1B3, and ATP2B1) and voltage-gated Na^+^ (SCN1A, SCN1B, and SCN2B) and K^+^ channels (KCNAB1, KCNAB2, KCNA2, KCNA3, and KCNH1) that regulate action potential firing patterns and intrinsic excitability. In addition, subunits for glutamate (GRIA2, GRIA3, GRID1, GRIK1, and GRIN1) and GABA (GABRA3 and GABRB2) receptors also differed in expression between control and AUD cases. Together, these findings suggest that AUD shifts intrinsic excitability and excitatory/inhibitory balance in OFC neurons, which is supported by preclinical evidence from alcohol dependent mice and heavy drinking monkeys that identified robust changes in cellular firing, synaptic plasticity, spine morphology, and pre- and post-synaptic glutamatergic function^10^. While we did not find evidence for proteomic changes in GABAergic signaling in the drinking macaques (life span of 25-30 years)^22^, their history with alcohol drinking was relatively short (6 months) compared with 30+ years of drinking in the AUD cases used in the current manuscript. In comparison, adaptations in presynaptic signaling proteins that regulate glutamatergic release were identified in both human AUD cases and heavy drinking cynomolgus macaques^22^, which is congruent with evidence showing increased frequency of spontaneous postsynaptic currents in the drinking monkeys. Thus, the proteomic results identified possible mechanisms underlying cross-species adaptations in cell firing and synaptic function of OFC neurons and demonstrated similar adaptations in both males and females with AUD.

### Predicting AUD

Using supervised machine learning for classification and prediction of AUD, we identified distinct functional classes of proteins that separate control from AUD cases. The first class of predictor proteins is known to regulate presynaptic vesicle cycling and exocytosis and includes SV2A, RIMS1, RAB3A, CADPS, and STXBP5. Eighteen of the predictor proteins are expressed in vesicular membranes, 9 of which are present in clathrin-coated membranes. This set of proteins that regulate clathrin-mediated endocytosis, including AP2A1 and AP2S1, which are large and small chains within the heterotetrameric AP-2 adaptor complex, respectively. Interestingly, these two proteins were also predictors of heavy alcohol drinking in our monkey OFC proteomics study^22^, and a polymorphism in *AP2A1* showed suggestive association with ‘MaxDrinks’^49^, a heritable trait and risk factor for developing AUD^50-52^. Bioinformatics also revealed that *AP2A1* is altered in postmortem brain in AUD^66^ and after alcohol exposure in mice^41, 48, 67^. *Ap2a1* is also present in QTLs for alcohol-related phenotypes (e.g., alcohol preference) and its expression correlated with voluntary drinking in BXDs. Furthermore, inbred strains that inherited the *Ap2a1 B* allele consumed high amounts of alcohol, supporting a cross-species role for AP2A1 in regulating excessive alcohol drinking.

A second major protein class that separate control from AUD cases regulate catabolic and metabolic processes within mitochondria. Additionally, mitochondria proteins involved in catabolic and metabolic processes also separated cases based on the severity of AUD. As chronic alcohol is known to cause alterations in presynaptic excitation-secretion coupling^68^, our results suggest that presynaptic adaptations in the OFC may be a core feature of AUD, whereas mitochondrial dysfunction could be present in individuals with a severe diagnosis of AUD. While some mechanisms of alcohol effects on presynapses are beginning to emerge, future work must determine if presynaptic proteins, such as those within the AP-2 complex, control excessive alcohol drinking or if pharmacological targeting of presynaptic proteins can reduce consumption and high rates of relapse in individuals with AUD. Likewise, further investigations into restoration of mitochondrial function are necessary to determine the potential as adjunct pharmacotherapy to improve cognitive control and treat relapse.

### AUD and social behaviors

Another intriguing finding from the current study was that OFC DEPs are associated with control of social behaviors, including social interactions and nonverbal communicative behavior. While the OFC is not widely studied in the context of social behaviors, there is cross-species evidence supporting a role of the OFC in regulating social interactions. Patients with OFC lesions show inappropriate social behavior and difficulty using facial expressions to guide social decisions^69, 70^. Studies in non-human primates reported that OFC neurons encode value of rewards obtained in social contexts and track social preferences and rank^71, 72^. In rodents, social interaction activates OFC neurons and 5-HT release^73^, and optogenetic manipulation of OFC projections to the basolateral amygdala alters social behavior^74, 75^. There is strong evidence that individuals with AUD have difficulties with social cognition, social decision-making, and emotion processing^69, 76^, which is proposed to contribute to the development and maintenance of excessive alcohol drinking and the increased risk for relapse^69^. While it is unclear if the DEPs represent innate biological differences or are a consequence of heavy drinking, clinical studies suggest that severity of deficits in facial emotion recognition tasks are related to greater lifetime drinking and longer durations of excessive drinking episodes, as well as poorer treatment outcome^77-79^. There is some evidence demonstrating social cognitive impairments in individuals with a family history of AUD^80-82^, and studies in high-risk individuals reported reduced OFC cortical thickness and volume^83, 84^. Of the alcohol-sensitive proteins linked to social behaviors, *SYT1, TSC1, SLC6A1, UPF3B, GRIA3, TSC2*, and *SCN1A* are high-risk genes for neurodevelopmental disorders with altered social skills, such as syndromic autism spectrum disorder (ASD). Moreover, *GPHN, GRIN1, OPHN1*, and *PC* are genes associated with high risk for developing non-syndromic ASD. Consistent with a change in expression due to excessive drinking, our machine learning analysis identified two alcohol-sensitive proteins (GRIN1 and AP3B2) involved in social behaviors that separated AUD cases based on DSM-5 diagnosis severity. Thus, by identifying novel targets in the human OFC, these results highlight the need to determine a mechanistic role of the alcohol-sensitive proteins in controlling excessive drinking and deficits in social cognition in an effort to reduce relapse rates.

### OFC sexual dimorphism

Corroborating known sexual dimorphism in human brain structure^85, 86^, the current study identified differences in OFC proteins that regulate ECM structure and signaling, as well as axonogenesis, between males and females. There were 9 ECM structural proteins that differed by sex, including COL4A1, COL4A2, HSPG2, LAMA2, LAMB2, and LAMC1, which are key components of basement membranes. This finding is consistent with a study using mouse cortex that reported sex differences in laminin and collagen ECM genes *Lama4, Lama5*, and *Col4a1*^87^. While LAMA5 protein expression did not differ in our study (and LAMA4 was not detected), COL4A1 expression was reduced in male brains in both species. Comparing males to females, we also observed reduced expression of laminin α, β, and γ, which are necessary for the assembly of the cruciform structure of heterotrimeric laminins that support ECM structure. Human OFC gray matter volume and cortical ribbon complexity are different in males and females^85, 88^, as is the cingulum axonal bundle that contains OFC afferents^89^. Brain viscoelasticity, which depends on ECM and axonal structure^90^, also shows sex differences in human^91-93^ and mouse^87^ cortex. In both species, male brains were softer than female brains. To our knowledge, this is the first evidence to identify molecular differences in ECM and axonal proteins that could explain sexual dimorphism in the structure of both white and gray matter in the human OFC. Some of these ECM proteins (TINAGL1, COL4A1, COL4A2, VWA1) were also differentially expressed in AUD cases, consistent with adaptations in the ECM by alcohol and other abused substances^48, 94-97^.

## Conclusion

Findings from our unbiased analysis of the OFC proteome in AUD cases identified adaptations in synaptic and mitochondrial proteins that predicted AUD and its severity. The results are consistent with functional adaptations reported in the OFC of alcohol dependent mice and heavy drinking non-human primates, demonstrating convergence of alcohol effects in mice, monkeys, and humans. Moreover, we also identified alcohol-sensitive OFC proteins that could drive alcohol drinking and social dysfunction in AUD and proteins that support ECM structure that explain sex-specific neuroanatomical differences in the OFC.

## ACKNOWLEDGMENTS

Tissues were received from the New South Wales Brain Tissue Resource Centre at the University of Sydney which is supported by the University of Sydney. Research reported in this publication was supported by the National Institute of Alcohol Abuse and Alcoholism of the National Institutes of Health under Award Number R28AA012725 The content is solely the responsibility of the authors and does not represent the official views of the National Institutes of Health. NIH grants to PJM (R01 AA023288) and Charleston Alcohol Research Center to PJM and JJW (P50 AA010761) supported this work. Data were acquired by LEB (MUSC Mass Spectrometry Facility) with support from NIH grants (S10 OD025126 and P30GM140964). Proteomic data analysis was performed by PW (OHSU Proteomics Shared Resource) with partial support from NIH core grants P30EY010572 and P30CA069533.

## AUTHOR CONTRIBUTIONS

P.J.M. and J.J.W. conceived and coordinated the project. L.E.B. designed and conducted the TMTpro experiment. P.A.W. processed the proteomics data. S.B. and P.J.M. analyzed the data. P.J.M. performed the concordance analysis and bioinformatics and reverse genetics experiments. C.M. performed the machine learning analysis. P.J.M. wrote the manuscript. P.J.M, J.J.W., L.E.B., S.B., P.A.W., and C.M. revised and edited the manuscript. P.J.M. supervised the overall project and provided funding and resources.

## INCLUSION AND DIVERSITY

We support inclusive, diverse, and equitable conduct of research.

## SUPPLEMENTAL FIGURE

**Supplemental Figure 1.**
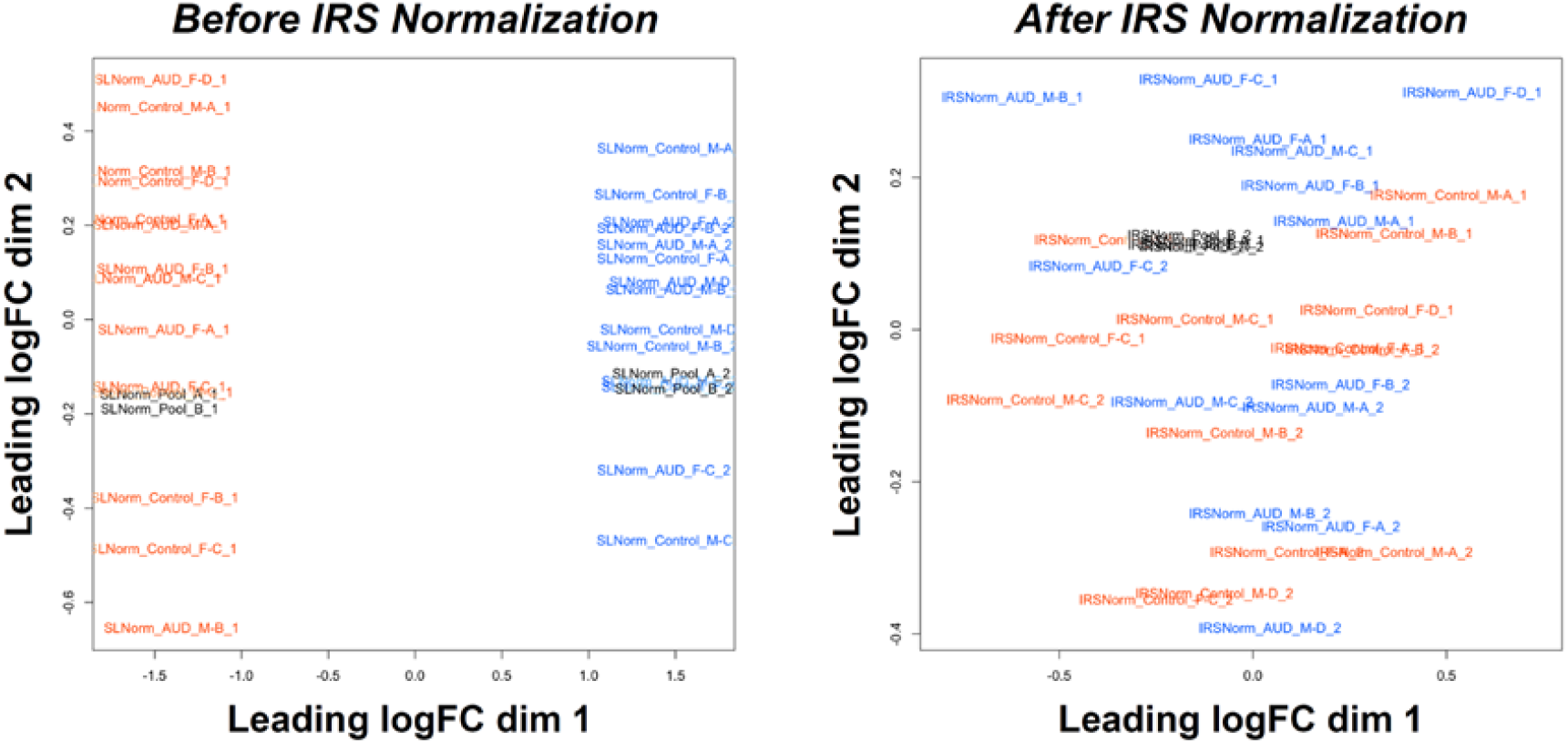
IKS normalization ot two IM I plexes. IKS normalizes batch effects across two 16-plex TMT kits processed in separate MS/MS runs. Left panel’. Multi-dimensional scaling (MDS) plot demonstrating batch-like effects of reporter ion intensities. In the MDS plot, all samples from TMT kit 1 (orange font) localize to opposite dimensions compared with samples in TMT kit 2 (blue font) prior to IRS normalization. Right panel: IRS removes batch effects from different MS/MS runs. The pooled reference channels are shown in black font, and the pooled samples for the individual kits locate to opposite dimensions before IRS normalization. As expected, all four pooled samples localize to near-exact dimensions in the MDS plot following IRS normalization.

**Supplemental Figure 2.**
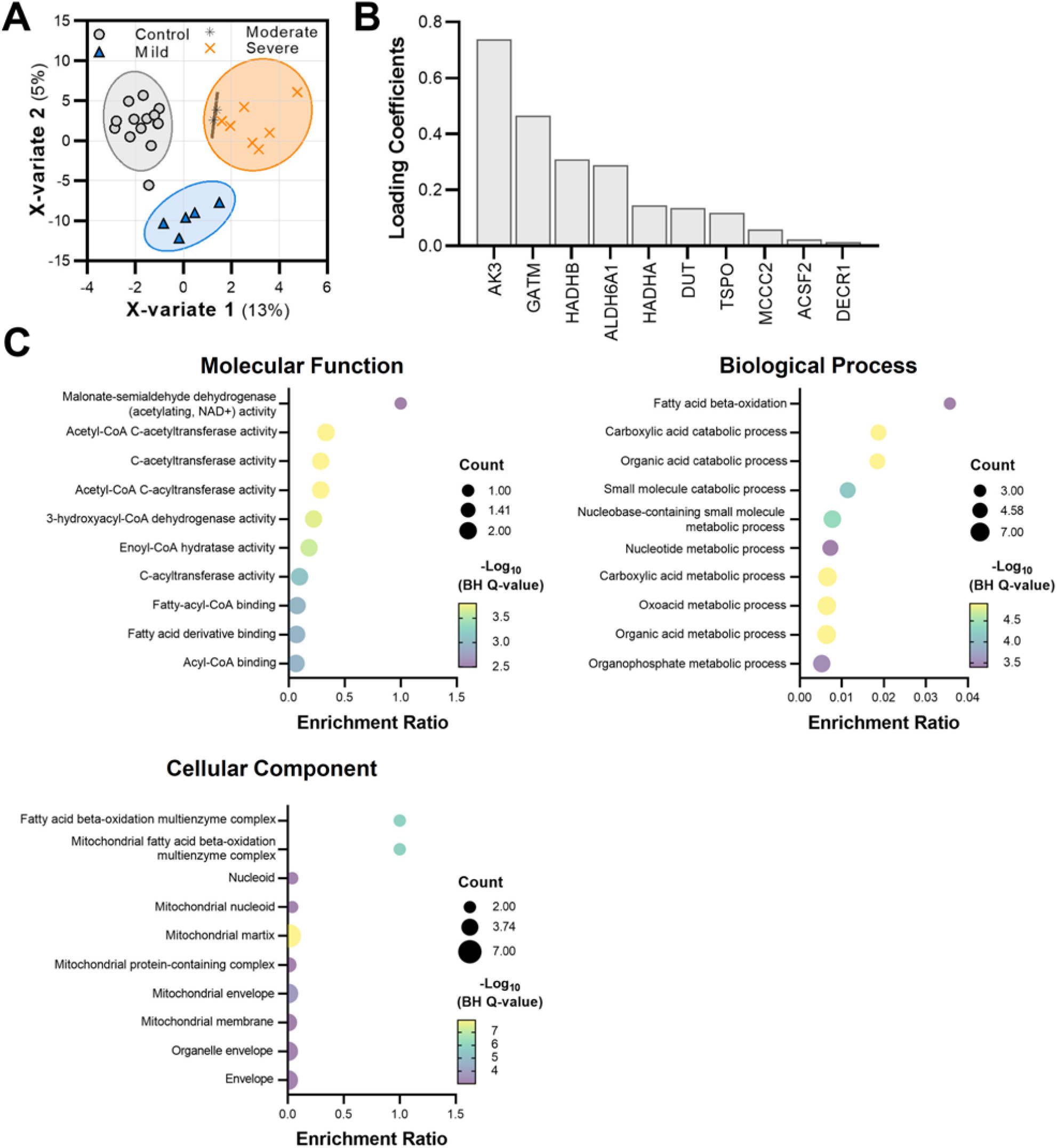
Identification of proteins that separate control and severity of AUD cases (mild, moderate, and severe AUD). **A** sPLS-DA performed on all proteins separated cases from controls and AUD based on severity of diagnosis. **B** Loading coefficients for the first component of the sPLS-DA model. Bar length indicated importance of each protein to the model. **C** Gene ontology enrichment analysis for the 10 proteins that separate control and the severity of AUD diagnosis.

## References

1. Collaborators GBDRF. Global burden of 87 risk factors in 204 countries and territories, 1990-2019: a systematic analysis for the Global Burden of Disease Study 2019. Lancet 2020; 396(10258): 1223–1249.

2. Spectrum N. Alcohol-related deaths continued to increase in 2021. vol. 15. National Institute of Alcohol Abuse and Alcoholism 2023.

3. Glantz MD, Bharat C, Degenhardt L, Sampson NA, Scott KM, Lim CCW et al. The epidemiology of alcohol use disorders cross-nationally: Findings from the World Mental Health Surveys. Addict Behav 2020; 102: 106128.

4. DSM-5. Diagnostic and statistical manual of mental disorders : DSM-5-TR. 5th ed. Text Revision DSM-5-TR edn. American Psychiatric Association: Arlington, VA, 2022.

5. Koob GF. Anhedonia, Hyperkatifeia, and Negative Reinforcement in Substance Use Disorders. Curr Top Behav Neurosci 2022.

6. Badanich K, Mulholland P, Beckley J, Trantham-Davidson H, Woodward J. Ethanol reduces neuronal excitability of lateral orbitofrontal cortex neurons via a glycine receptor dependent mechanism. Neuropsycopharmacology 2013; 38(7): 1176–1188.

7. Nimitvilai S, Lopez MF, Mulholland PJ, Woodward JJ. Chronic Intermittent Ethanol Exposure Enhances the Excitability and Synaptic Plasticity of Lateral Orbitofrontal Cortex Neurons and Induces a Tolerance to the Acute Inhibitory Actions of Ethanol. Neuropsychopharmacology 2016; 41(4): 1112–1127.

8. Cannady R, Nimitvilai-Roberts S, Jennings SD, Woodward JJ, Mulholland PJ. Distinct Region- and Time-Dependent Functional Cortical Adaptations in C57BL/6J Mice after Short and Prolonged Alcohol Drinking. eNeuro 2020; 7(3).

9. Moorman DE. The role of the orbitofrontal cortex in alcohol use, abuse, and dependence. Prog Neuropsychopharmacol Biol Psychiatry 2018; 87(Pt A): 85–107.

10. Shields CN, Gremel CM. Review of Orbitofrontal Cortex in Alcohol Dependence: A Disrupted Cognitive Map? Alcoholism, clinical and experimental research 2020; 44(10): 1952–1964.

11. Hernandez JS, Moorman DE. Orbitofrontal Cortex Encodes Preference for Alcohol. eNeuro 2020; 7(4).

12. Jokisch D, Roser P, Juckel G, Daum I, Bellebaum C. Impairments in learning by monetary rewards and alcohol-associated rewards in detoxified alcoholic patients. Alcoholism, clinical and experimental research 2014; 38(7): 1947–1954.

13. Badanich KA, Fakih ME, Gurina TS, Roy EK, Hoffman JL, Uruena-Agnes AR et al. Reversal learning and experimenter-administered chronic intermittent ethanol exposure in male rats. Psychopharmacology 2016.

14. Badanich K, Becker H, Woodward J. Effects of chronic intermittent ethanol exposure on orbitofrontal and medial prefrontal cortex-dependent behaviors in mice. Behavioral neuroscience 2011; 125(6): 879–891.

15. McMurray MS, Amodeo LR, Roitman JD. Effects of voluntary alcohol intake on risk preference and behavioral flexibility during rat adolescence. PLoS One 2014; 9(7): e100697.

16. Jedema HP, Carter MD, Dugan BP, Gurnsey K, Olsen AS, Bradberry CW. The acute impact of ethanol on cognitive performance in rhesus macaques. Cerebral cortex 2011; 21(8): 1783–1791.

17. Renteria R, Baltz ET, Gremel CM. Chronic alcohol exposure disrupts top-down control over basal ganglia action selection to produce habits. Nat Commun 2018; 9(1): 211.

18. Volkow ND, Wang GJ, Overall JE, Hitzemann R, Fowler JS, Pappas N et al. Regional brain metabolic response to lorazepam in alcoholics during early and late alcohol detoxification. Alcoholism, clinical and experimental research 1997; 21(7): 1278–1284.

19. Myrick H, Anton RF, Li X, Henderson S, Drobes D, Voronin K et al. Differential brain activity in alcoholics and social drinkers to alcohol cues: relationship to craving. Neuropsychopharmacology : official publication of the American College of Neuropsychopharmacology 2004; 29(2): 393–402.

20. Schacht JP, Yeongbin I, Hoffman M, Voronin KE, Book SW, Anton RF. Effects of pharmacological and genetic regulation of COMT activity in alcohol use disorder: a randomized, placebo-controlled trial of tolcapone. Neuropsychopharmacology : official publication of the American College of Neuropsychopharmacology 2022.

21. Gioia DA, Woodward JJ. Altered Activity of Lateral Orbitofrontal Cortex Neurons in Mice following Chronic Intermittent Ethanol Exposure. eNeuro 2021; 8(2).

22. Nimitvilai S, Uys JD, Woodward JJ, Randall PK, Ball LE, Williams RW et al. Orbitofrontal Neuroadaptations and Cross-Species Synaptic Biomarkers in Heavy-Drinking Macaques. J Neurosci 2017; 37(13): 3646–3660.

23. Ongur D, Ferry AT, Price JL. Architectonic subdivision of the human orbital and medial prefrontal cortex. The Journal of comparative neurology 2003; 460(3): 425–449.

24. Li J, Van Vranken JG, Pontano Vaites L, Schweppe DK, Huttlin EL, Etienne C et al. TMTpro reagents: a set of isobaric labeling mass tags enables simultaneous proteome-wide measurements across 16 samples. Nature methods 2020; 17(4): 399–404.

25. McAlister GC, Nusinow DP, Jedrychowski MP, Wuhr M, Huttlin EL, Erickson BK et al. MultiNotch MS3 enables accurate, sensitive, and multiplexed detection of differential expression across cancer cell line proteomes. Analytical chemistry 2014; 86(14): 7150–7158.

26. Eng JK, Jahan TA, Hoopmann MR. Comet: an open-source MS/MS sequence database search tool. Proteomics 2013; 13(1): 22–24.

27. Wilmarth PA, Riviere MA, David LL. Techniques for accurate protein identification in shotgun proteomic studies of human, mouse, bovine, and chicken lenses. J Ocul Biol Dis Infor 2009; 2(4): 223–234.

28. Chambers MC, Maclean B, Burke R, Amodei D, Ruderman DL, Neumann S et al. A cross-platform toolkit for mass spectrometry and proteomics. Nat Biotechnol 2012; 30(10): 918–920.

29. McDonald WH, Tabb DL, Sadygov RG, MacCoss MJ, Venable J, Graumann J et al. MS1, MS2, and SQT-three unified, compact, and easily parsed file formats for the storage of shotgun proteomic spectra and identifications. Rapid Commun Mass Spectrom 2004; 18(18): 2162–2168.

30. Keller A, Nesvizhskii AI, Kolker E, Aebersold R. Empirical statistical model to estimate the accuracy of peptide identifications made by MS/MS and database search. Analytical chemistry 2002; 74(20): 5383–5392.

31. Elias JE, Gygi SP. Target-decoy search strategy for increased confidence in large-scale protein identifications by mass spectrometry. Nature methods 2007; 4(3): 207–214.

32. Plubell DL, Wilmarth PA, Zhao Y, Fenton AM, Minnier J, Reddy AP et al. Extended Multiplexing of Tandem Mass Tags (TMT) Labeling Reveals Age and High Fat Diet Specific Proteome Changes in Mouse Epididymal Adipose Tissue. Molecular & cellular proteomics : MCP 2017; 16(5): 873–890.

33. Perez-Riverol Y, Bai J, Bandla C, Garcia-Seisdedos D, Hewapathirana S, Kamatchinathan S et al. The PRIDE database resources in 2022: a hub for mass spectrometry-based proteomics evidences. Nucleic Acids Res 2022; 50(D1): D543–D552.

34. Le Cao KA, Boitard S, Besse P. Sparse PLS discriminant analysis: biologically relevant feature selection and graphical displays for multiclass problems. BMC Bioinformatics 2011; 12: 253.

35. McGuier NS, Rinker JA, Cannady R, Fulmer DB, Jones SR, Hoffman M et al. Identification and validation of midbrain Kcnq4 regulation of heavy alcohol consumption in rodents. Neuropharmacology 2018; 138: 10–19.

36. Padula AE, Griffin WC, 3rd, Lopez MF, Nimitvilai S, Cannady R, McGuier NS et al. KCNN Genes that Encode Small-Conductance Ca2+-Activated K+ Channels Influence Alcohol and Drug Addiction. Neuropsychopharmacology 2015; 40(8): 1928–1939.

37. Padula AE, Rinker JA, Lopez MF, Mulligan MK, Williams RW, Becker HC et al. Bioinformatics identification and pharmacological validation of Kcnn3/K(Ca)2 channels as a mediator of negative affective behaviors and excessive alcohol drinking in mice. Translational psychiatry 2020; 10(1): 414.

38. Rinker JA, Fulmer DB, Trantham-Davidson H, Smith ML, Williams RW, Lopez MF et al. Differential potassium channel gene regulation in BXD mice reveals novel targets for pharmacogenetic therapies to reduce heavy alcohol drinking. Alcohol 2017; 58: 33–45.

39. McGuier NS, Griffin WC, 3rd, Gass JT, Padula AE, Chesler EJ, Mulholland PJ. Kv7 channels in the nucleus accumbens are altered by chronic drinking and are targets for reducing alcohol consumption. Addiction biology 2015.

40. Baker EJ, Jay JJ, Bubier JA, Langston MA, Chesler EJ. GeneWeaver: a web-based system for integrative functional genomics. Nucleic acids research 2012; 40(Database issue): D1067–1076.

41. Philip VM, Duvvuru S, Gomero B, Ansah TA, Blaha CD, Cook MN et al. High-throughput behavioral phenotyping in the expanded panel of BXD recombinant inbred strains. Genes Brain Behav 2010; 9(2): 129–159.

42. Dickson PE, Miller MM, Calton MA, Bubier JA, Cook MN, Goldowitz D et al. Systems genetics of intravenous cocaine self-administration in the BXD recombinant inbred mouse panel. Psychopharmacology 2016; 233(4): 701–714.

43. Lopez MF, Miles MF, Williams RW, Becker HC. Variable effects of chronic intermittent ethanol exposure on ethanol drinking in a genetically diverse mouse cohort. Alcohol 2017; 58: 73–82.

44. Faul F, Erdfelder E, Lang AG, Buchner A. G*Power 3: a flexible statistical power analysis program for the social, behavioral, and biomedical sciences. Behav Res Methods 2007; 39(2): 175–191.

45. Leek JT, Johnson WE, Parker HS, Jaffe AE, Storey JD. The sva package for removing batch effects and other unwanted variation in high-throughput experiments. Bioinformatics 2012; 28(6): 882–883.

46. Chen J, Bardes EE, Aronow BJ, Jegga AG. ToppGene Suite for gene list enrichment analysis and candidate gene prioritization. Nucleic Acids Res 2009; 37(Web Server issue): W305–311.

47. Cahill KM, Huo Z, Tseng GC, Logan RW, Seney ML. Improved identification of concordant and discordant gene expression signatures using an updated rank-rank hypergeometric overlap approach. Sci Rep 2018; 8(1): 9588.

48. Uys JD, McGuier NS, Gass JT, Griffin WC, 3rd, Ball LE, Mulholland PJ. Chronic intermittent ethanol exposure and withdrawal leads to adaptations in nucleus accumbens core postsynaptic density proteome and dendritic spines. Addict Biol 2016; 21(3): 560–574.

49. Kapoor M, Wang JC, Wetherill L, Le N, Bertelsen S, Hinrichs AL et al. A meta-analysis of two genome-wide association studies to identify novel loci for maximum number of alcoholic drinks. Hum Genet 2013; 132(10): 1141–1151.

50. Grant JD, Agrawal A, Bucholz KK, Madden PA, Pergadia ML, Nelson EC et al. Alcohol consumption indices of genetic risk for alcohol dependence. Biological psychiatry 2009; 66(8): 795–800.

51. Saccone NL, Kwon JM, Corbett J, Goate A, Rochberg N, Edenberg HJ et al. A genome screen of maximum number of drinks as an alcoholism phenotype. Am J Med Genet 2000; 96(5): 632–637.

52. Schuckit MA, Smith TL, Danko GP, Bucholz KK, Agrawal A, Dick DM et al. Predictors of subgroups based on maximum drinks per occasion over six years for 833 adolescents and young adults in COGA. J Stud Alcohol Drugs 2014; 75(1): 24–34.

53. Wang X, Pandey AK, Mulligan MK, Williams EG, Mozhui K, Li Z et al. Joint mouse-human phenome-wide association to test gene function and disease risk. Nat Commun 2016; 7: 10464.

54. Caruana NJ, Stroud DA. The road to the structure of the mitochondrial respiratory chain supercomplex. Biochem Soc Trans 2020; 48(2): 621–629.

55. Qin L, Vetreno RP, Crews FT. NADPH oxidase and endoplasmic reticulum stress is associated with neuronal degeneration in orbitofrontal cortex of individuals with alcohol use disorder. Addiction biology 2023; 28(1): e13262.

56. Flatscher-Bader T, van der Brug M, Hwang JW, Gochee PA, Matsumoto I, Niwa S et al. Alcohol-responsive genes in the frontal cortex and nucleus accumbens of human alcoholics. Journal of neurochemistry 2005; 93(2): 359–370.

57. Sokolov BP, Jiang L, Trivedi NS, Aston C. Transcription profiling reveals mitochondrial, ubiquitin and signaling systems abnormalities in postmortem brains from subjects with a history of alcohol abuse or dependence. J Neurosci Res 2003; 72(6): 756–767.

58. Liu J, Lewohl JM, Dodd PR, Randall PK, Harris RA, Mayfield RD. Gene expression profiling of individual cases reveals consistent transcriptional changes in alcoholic human brain. Journal of neurochemistry 2004; 90(5): 1050–1058.

59. Shang P, Lindberg D, Starski P, Peyton L, Hong SI, Choi S et al. Chronic Alcohol Exposure Induces Aberrant Mitochondrial Morphology and Inhibits Respiratory Capacity in the Medial Prefrontal Cortex of Mice. Front Neurosci 2020; 14: 561173.

60. Jung ME, Metzger DB. Aberrant histone acetylation promotes mitochondrial respiratory suppression in the brain of alcoholic rats. J Pharmacol Exp Ther 2015; 352(2): 258–266.

61. Reddy VD, Padmavathi P, Kavitha G, Saradamma B, Varadacharyulu N. Alcohol-induced oxidative/nitrosative stress alters brain mitochondrial membrane properties. Mol Cell Biochem 2013; 375(1-2): 39–47.

62. Vetreno RP, Qin L, Coleman LG, Jr., Crews FT. Increased Toll-like Receptor-MyD88-NFkappaB-Proinflammatory neuroimmune signaling in the orbitofrontal cortex of humans with alcohol use disorder. Alcoholism, clinical and experimental research 2021; 45(9): 1747–1761.

63. Miguel-Hidalgo JJ, Overholser JC, Meltzer HY, Stockmeier CA, Rajkowska G. Reduced glial and neuronal packing density in the orbitofrontal cortex in alcohol dependence and its relationship with suicide and duration of alcohol dependence. Alcoholism, clinical and experimental research 2006; 30(11): 1845–1855.

64. Ruggiero A, Katsenelson M, Slutsky I. Mitochondria: new players in homeostatic regulation of firing rate set points. Trends Neurosci 2021; 44(8): 605–618.

65. Styr B, Gonen N, Zarhin D, Ruggiero A, Atsmon R, Gazit N et al. Mitochondrial Regulation of the Hippocampal Firing Rate Set Point and Seizure Susceptibility. Neuron 2019; 102(5): 1009–1024 e1008.

66. Ponomarev I, Wang S, Zhang L, Harris RA, Mayfield RD. Gene coexpression networks in human brain identify epigenetic modifications in alcohol dependence. J Neurosci 2012; 32(5): 1884–1897.

67. Wolen AR, Phillips CA, Langston MA, Putman AH, Vorster PJ, Bruce NA et al. Genetic dissection of acute ethanol responsive gene networks in prefrontal cortex: functional and mechanistic implications. PloS one 2012; 7(4): e33575.

68. Lovinger DM. Presynaptic Ethanol Actions: Potential Roles in Ethanol Seeking. Handb Exp Pharmacol 2018; 248: 29–54.

69. Le Berre AP. Emotional processing and social cognition in alcohol use disorder. Neuropsychology 2019; 33(6): 808–821.

70. Willis ML, Palermo R, Burke D, McGrillen K, Miller L. Orbitofrontal cortex lesions result in abnormal social judgements to emotional faces. Neuropsychologia 2010; 48(7): 2182–2187.

71. Watson KK, Platt ML. Social signals in primate orbitofrontal cortex. Curr Biol 2012; 22(23): 2268–2273.

72. Azzi JC, Sirigu A, Duhamel JR. Modulation of value representation by social context in the primate orbitofrontal cortex. Proceedings of the National Academy of Sciences of the United States of America 2012; 109(6): 2126–2131.

73. Unger EK, Keller JP, Altermatt M, Liang R, Matsui A, Dong C et al. Directed Evolution of a Selective and Sensitive Serotonin Sensor via Machine Learning. Cell 2020; 183(7): 1986–2002 e1926.

74. Kuniishi H, Nakatake Y, Sekiguchi M, Yamada M. Adolescent social isolation induces distinct changes in the medial and lateral OFC-BLA synapse and social and emotional alterations in adult mice. Neuropsychopharmacology : official publication of the American College of Neuropsychopharmacology 2022; 47(9): 1597–1607.

75. Jennings JH, Kim CK, Marshel JH, Raffiee M, Ye L, Quirin S et al. Interacting neural ensembles in orbitofrontal cortex for social and feeding behaviour. Nature 2019; 565(7741): 645–649.

76. Gautier M, Pabst A, Maurage P. Social decision making in severe alcohol use disorder: Scoping review and experimental perspectives. Alcoholism, clinical and experimental research 2021; 45(8): 1548–1559.

77. Charlet K, Schlagenhauf F, Richter A, Naundorf K, Dornhof L, Weinfurtner CE et al. Neural activation during processing of aversive faces predicts treatment outcome in alcoholism. Addiction biology 2014; 19(3): 439–451.

78. Bora E, Zorlu N. Social cognition in alcohol use disorder: a meta-analysis. Addiction 2017; 112(1): 40–48.

79. Valmas MM, Mosher Ruiz S, Gansler DA, Sawyer KS, Oscar-Berman M. Social cognition deficits and associations with drinking history in alcoholic men and women. Alcoholism, clinical and experimental research 2014; 38(12): 2998–3007.

80. Schmidt T, Roser P, Ze O, Juckel G, Suchan B, Thoma P. Cortical thickness and trait empathy in patients and people at high risk for alcohol use disorders. Psychopharmacology 2017; 234(23-24): 3521–3533.

81. Hulvershorn LA, Finn P, Hummer TA, Leibenluft E, Ball B, Gichina V et al. Cortical activation deficits during facial emotion processing in youth at high risk for the development of substance use disorders. Drug Alcohol Depend 2013; 131(3): 230–237.

82. Hill SY, Kostelnik B, Holmes B, Goradia D, McDermott M, Diwadkar V et al. fMRI BOLD response to the eyes task in offspring from multiplex alcohol dependence families. Alcoholism, clinical and experimental research 2007; 31(12): 2028–2035.

83. Hill SY, Wellman JL, Zezza N, Steinhauer SR, Sharma V, Holmes B. Epigenetic Effects in HPA Axis Genes Associated with Cortical Thickness, ERP Components and SUD Outcome. Behav Sci (Basel) 2022; 12(10).

84. Hill SY, Wang S, Kostelnik B, Carter H, Holmes B, McDermott M et al. Disruption of orbitofrontal cortex laterality in offspring from multiplex alcohol dependence families. Biological psychiatry 2009; 65(2): 129–136.

85. Xin J, Zhang Y, Tang Y, Yang Y. Brain Differences Between Men and Women: Evidence From Deep Learning. Front Neurosci 2019; 13: 185.

86. Ritchie SJ, Cox SR, Shen X, Lombardo MV, Reus LM, Alloza C et al. Sex Differences in the Adult Human Brain: Evidence from 5216 UK Biobank Participants. Cerebral cortex 2018; 28(8): 2959–2975.

87. Batzdorf CS, Morr AS, Bertalan G, Sack I, Silva RV, Infante-Duarte C. Sexual Dimorphism in Extracellular Matrix Composition and Viscoelasticity of the Healthy and Inflamed Mouse Brain. Biology (Basel) 2022; 11(2).

88. Nazlee N, Waiter GD, Sandu AL. Age-associated sex and asymmetry differentiation in hemispheric and lobar cortical ribbon complexity across adulthood: A UK Biobank imaging study. Hum Brain Mapp 2022; 44(1): 49–65.

89. Heilbronner SR, Haber SN. Frontal cortical and subcortical projections provide a basis for segmenting the cingulum bundle: implications for neuroimaging and psychiatric disorders. The Journal of neuroscience : the official journal of the Society for Neuroscience 2014; 34(30): 10041–10054.

90. Guo J, Bertalan G, Meierhofer D, Klein C, Schreyer S, Steiner B et al. Brain maturation is associated with increasing tissue stiffness and decreasing tissue fluidity. Acta Biomater 2019; 99: 433–442.

91. Arani A, Murphy MC, Glaser KJ, Manduca A, Lake DS, Kruse SA et al. Measuring the effects of aging and sex on regional brain stiffness with MR elastography in healthy older adults. Neuroimage 2015; 111: 59–64.

92. Sack I, Beierbach B, Wuerfel J, Klatt D, Hamhaber U, Papazoglou S et al. The impact of aging and gender on brain viscoelasticity. Neuroimage 2009; 46(3): 652–657.

93. Wuerfel J, Paul F, Beierbach B, Hamhaber U, Klatt D, Papazoglou S et al. MR-elastography reveals degradation of tissue integrity in multiple sclerosis. Neuroimage 2010; 49(3): 2520–2525.

94. Mulholland PJ, Chandler LJ, Kalivas PW. Signals from the Fourth Dimension Regulate Drug Relapse. Trends Neurosci 2016; 39(7): 472–485.

95. Lasek AW. Effects of Ethanol on Brain Extracellular Matrix: Implications for Alcohol Use Disorder. Alcoholism, clinical and experimental research 2016; 40(10): 2030–2042.

96. Kruyer A, Chioma VC, Kalivas PW. The Opioid-Addicted Tetrapartite Synapse. Biological psychiatry 2020; 87(1): 34–43.

97. Seney ML, Kim SM, Glausier JR, Hildebrand MA, Xue X, Zong W et al. Transcriptional Alterations in Dorsolateral Prefrontal Cortex and Nucleus Accumbens Implicate Neuroinflammation and Synaptic Remodeling in Opioid Use Disorder. Biological psychiatry 2021; 90(8): 550–562.

98. Sheedy D, Garrick T, Dedova I, Hunt C, Miller R, Sundqvist N et al. An Australian Brain Bank: a critical investment with a high return! Cell and tissue banking 2008; 9(3): 205–216.

